# Variable penetrance of schizophrenia in 22q11.2 deletion syndrome is associated with impaired, but reversible, neuronal mitophagy

**DOI:** 10.64898/2026.11.13.766189

**Authors:** Eleonora Stronati, Adam J. Rossano, Jianping. Li, Amelia. Stoesser, Douglas C. Wallace, Cecilia Giulivi, Stewart A. Anderson

**Affiliations:** Department of Child and Adolescent Psychiatry, Children’s Hospital of Philadelphia, Philadelphia, PA 19104, USA; Department of Psychiatry, Perelman School of Medicine, University of Pennsylvania, Philadelphia, PA 19104, USA; Department of Psychiatry, the First Affiliated Hospital, Zhejiang University School of Medicine, Hangzhou, 310003, China; Department of Molecular Biosciences, School of Veterinary Medicine, University of California Davis, Davis, CA 95616, USA; University of California Medical Investigations of Neurodevelopmental Disorders Institute (MIND Institute), University of California Health, Sacramento, CA 95817, USA

## Abstract

The 22q11.2 deletion syndrome (22qDS) is a major genetic risk factor for schizophrenia (Sz). We found that iPSC-derived neurons (iNeurons) from 22qDS +Sz individuals exhibit weaker mitochondrial energetics, whereas iNeurons from individuals with the same deletion but no psychosis (22qDS-Sz) upregulate expression of mitochondrial biogenesis-related genes. Here, we investigated why the +Sz group fails to undergo this compensation, finding that 22qDS +Sz iNeurons have older and more depolarized mitochondria than 22qDS - Sz or controls. Transcriptomic and proteomic analyses revealed downregulation of mitophagy-related pathways, and live imaging revealed elevated lysosomal pH. Restoring lysosomal acidity with acidifying nanoparticles improved mitophagy and, remarkably, increased the expression of mitochondrial biogenesis-related transcription factors in the +Sz neurons. We conclude that lysosomal dysfunction and impaired mitophagy differentiate 22qDS individuals who develop schizophrenia from those who do not. Mitophagy could be a novel therapeutic or preventative opportunity for Sz in the context of genetic neurodevelopmental disorders.

## Introduction

Schizophrenia (Sz) is a heterogeneous disorder generally characterized by adolescent onset of psychosis preceded by deteriorations of social and cognitive functioning(*1, 2*). Although genome-wide associated studies particularly implicate polygenic hits to synaptic function as contributors to the development of Sz(*3*), mitochondrial dysfunction has also been implicated(*4, 5*). For example, gene expression alterations in neocortical pyramidal neurons suggest disruption of mitochondrial function(*6*), and other studies reported mitochondrial defects in lymphocytes as well as neural progenitors, cerebral organoids, and cortical interneurons, derived from induced pluripotent stem cells (iPSCs) from patients with Sz relative to healthy controls(*7–11*).

Sz is a highly heritable disorder in which genetic risk is primarily polygenic, but the most common single genetic risk factor for Sz is the 22q11.2 deletion syndrome (22qDS), which occurs in about 1:3000 births(*12*). Roughly 25% of people with 22qDS develop Sz, a roughly 25x increase in risk, and they do so in a manner highly similar to idiopathic Sz(*13, 14*). Remarkably, seven of the 45 heterozygously-deleted genes encode proteins that localize to mitochondria, and at least four others strongly influence mitochondrial function(*15, 16*). Unsurprisingly, because of the links between schizophrenia, mitochondrial hypofunction, and 22qDS, many studies involving humans and animals have connected mitochondrial dysfunction with schizophrenia in 22qDS(*17–21*). Our own group, using iPSC-derived excitatory neurons (iNeurons) as well as lymphoblastoid cell lines (LCLs), found reduced Complex I activity and other OXPHOS-related abnormalities in the 22qDS and Sz (22qDS +Sz) group relative to both 22qDS without Sz (22qDS-Sz) and to neurotypical control groups(*22, 23*). We also observed an elevated expression of genes associated with mitochondrial biogenesis and mitophagy in the 22qDS-Sz iNeurons, suggesting that the capacity to render the mitochondrial population “younger” (mitochondrial proteins have variable rates of turnover)(*24*) was influencing the penetrance of Sz in the context of 22qDS(*22*).

Despite these observations, it is unknown whether actual mitochondrial biogenesis and turnover dynamics differ between 22qDS-Sz and 22qDS +Sz neurons, and why the apparent compensatory mechanisms observed at the transcript level in 22qDS-Sz fail to occur in 22qDS individuals +Sz. In this study, using iPSC-derived excitatory neurons from controls, 22qDS +Sz, and 22qDS-Sz(*25*), we test whether mitochondrial biogenesis and turnover are selectively upregulated in the 22qDS-Sz group, and investigate the mechanisms underlying the apparent absence of this response in the 22qDS +Sz group. Our findings indicate that mitochondria in 22qDS-Sz neurons indeed undergo accelerated biogenesis and turnover relative to controls, whereas mitochondria in 22qDS +Sz neurons exhibit reduced biogenesis and turnover. These results identify a previously uncharacterized divergence in mitochondrial dynamic regulation that may help explain differential Sz risk within 22qDS. Moreover, 22qDS +Sz iNeurons are accumulating partially depolarized mitochondria due to a slowing of mitophagy, which appears to be due to elevated pH in the lysosomes in the 22qDS +Sz group. Correcting lysosomal pH with acidifying nanoparticles (AcNPs) improves mitochondrial defects in this group and increases mitochondrial turnover. These results have important clinical implications, as enhancers of mitophagy are being actively studied as novel therapeutics for neurodegenerative disorders(*26–28*). Since defects in mitophagy have also been implicated in some instances of Sz outside of the 22qDS context(*29–31*), these results could have therapeutic implications for some individuals in the broader population of people with or at high risk for schizophrenia.

## Material and Methods

### IPSC lines

IPSC lines were previously published and kindly provided by Dr. Sergiu Pasca, Stanford University(*25*), and most lines were previously used by the Anderson lab(*22*). Supplemental Table 1 includes additional information. Importantly, the +Sz and-Sz groups had similar mean IQs.

### Neuronal Differentiation of iPSCs

The iPSC lines were infected with lentiviral vectors using lentiblast (OZ Bioscience): TetO-mNgn2-T2A-PuroR and Ubiq-rTTA as previously described(*23*). Neuronal differentiation was performed using an established method of doxycycline induction x 10 days initially together with transient dual-smad inhibition(*32*). At the Neural Progenitor stage (NPCs), 1*10^6 cells are frozen in batches and used as necessary. Cells are plated on poly-ornithine/laminin coated glass coverslips at 80000 cells/well, previously plated with a carpet of rat astrocytes. NPCs are plated into Neurobasal medium (Gibco) with B27 and N2 (Life Technologies) supplemented with doxycycline (2ug/mL) until differentiation day (DD) 10. From DD24 iNeurons are maintained in BrainPhys (Gibco) for 7 days until further analysis.

### Primary rat glial cells culture

Isolated E17 Sprague Dawley rat cortex was obtained from the UPenn NRU Core. Cells were plated onto uncoated 10 cm dishes at 1.5 million cells/ml. Cells were grown in NM-15 media (Eagle’s MEM with Earle’s salts and 2 mM l-glutamine, 15% heat-inactivated fetal bovine serum, 6 mg/ml glucose, 0.5 U/ml penicillin, and 0.5 μg/ml streptomycin). Medium exchanges occurred every 5 days, and rat glia grew for at least 2 weeks. Before glial harvest, the glial cells were shaken at 37 °C for 3 h at 230 rpm to remove microglia. The glial cells were then dissociated with Accutase (Life Technologies), spun down for 5 min at 1000 rpm, resuspended in Neurobasal/B27 media, and plated onto the neurons at 100k cells/well of a 24-well plate.

### Mitochondrial DNA copy number

Genomic DNA extraction was performed using DNAGenomic (Zymo), following the manufacturer’s instructions. All reactions were performed in 384-well reaction plates (Applied Biosystems), using SYBR green master mix (Applied Biosystems) on a Quant Studio 7 device. For each reaction, 3 ng/µl of DNA and 3 µM of primers (listed in the table) were applied. Relative mtDNA copy number (mtDNA amount/nDNA amount) was calculated by a comparative Ct method, using the following equation: mtDNA/nDNA = 2−ΔCt. Primer used are listed in Table S2.

### Western Blotting

Cells were harvested and total cell extracts were prepared in cold RIPA Buffer Lysis (Millipore) and cleared by centrifugation for 10 min at 10,000 g at 4C. Protein concentration was determined by BCA protein assay (Thermo Scientific). 20ug of sample were separated on a 4-20% SDS-PAGE gel (Biorad) and transferred onto nitrocellulose membranes using a Ublot gel transfer device (Biorad). Membranes were blocked for 1 hour at room temperature in TBS blocking buffer (Licor). Primary antibodies (LAMP1(Abcam, ab108597, 1:1000), ATG4B (Abcam, 199537, 1:1000), LC3B (Cell Signaling, 2775S, 1:500), TOMM20 (BD Transduction, 612278, 1:400), OPTN (Invitrogen, 711879, 1:700), ACTB (Cell Signaling, 3700S, 1:5000) were incubated in blocking buffer overnight at 4C and then incubated for 1 hour at room temperature with secondary antibody in blocking buffer. The protein bands were visualized using the Odyssey Infrared Imaging System LI-COR Biosciences.

### Immunostaining

iNeurons were fixed for 20 min in 4% PFA-1% sucrose (PFA, Sigma, P6148; D-sucrose, Fisher Scientific, BP220-1) in PBS (150 mM NaCl, 50 mM NaPO4, pH 7.4). Cells were washed twice in PBS, and stored in PBS at 4°C until subsequent processing. On the day of imaging, cells were permeabilized for 5 min in 0.1% TX-100 (Thermo Fisher Scientific, BP151-100) in PBS, washed with PBS, and blocked for 45 min in PBS supplemented with 5% BSA (Sigma) and 0.02% Saponin (Sigma). Samples were labeled with primary antibodies (LAMP1 (DHSB H4A3, 1:500), MAP2 (synaptic System 188004 1:700) diluted in blocking solution overnight at 4°C. Cells were washed 3 times for 5 min each in PBS and then incubated in secondary antibody diluted in blocking solution for 1 hour at room temperature. DAPI was included in the secondary antibody incubation at 1µg/ml. Coverslips were then mounted in ProLong Gold (Thermo Fisher Scientific/Molecular Probes, P36930). Samples were imaged by a blinded observer on a Nikon Eclipse TiE. Images that would be quantitatively compared to each other used the same acquisition parameters across treatment conditions.

### Treatments and Live-cell imaging

iNeurons were transfected at DD28 and treated and analyzed after 72h, with the exception of mtMkGO that was analyzed after 5 days (DD33). The transfection of 500 ng/μl of construct was performed using Lipofectamine 2000 (Invitrogen) following manufacturer instructions. The treatments were imaged in artificial cerebrospinal fluid for a maximum of 30 min. Imaging solutions were designed to maintain ionic composition of BrainPhys(*33*) and contained 121.0mM NaCl, 4.2mM KCl, 1.1mM CaCl_2_, 1.0mM MgSO_4_, 29.0mM NaHCO_3_, 0.5mM Na_2_HPO_4_, 0.45mM Na_2_HPO_4_-H_2_0, 2.5mM Glucose, 0.5mM Na-pyruvate, and 5mM HEPES buffered to pH 7.4 with HCl and N-Methyl-D-glutamine (NMDG) as needed. Solutions with NaProp and NH4Cl were generated with equimolar replacement of NaCl. All solutions were continuously bubbled with 5% CO2 balanced with O2.

Live cell imaging was performed using a custom live multi-wavelength epifluorescent imaging system using Olympus LumPlan 40x or 60x water-immersion objectives. The epifluorescent system is built upon an Olympus BX51WI housing. An Excelitas Xcite NOVEM LED system was used for excitation wavelength switching with Chroma and Semrock filters and dichorics. Optical path for BFP imaging contained a 385nm LED with 378/52nm clean-up filter into a 414nm dichroic with a 460/60nm emission filter. Interleaved GFP/RFP fluorescence (for either mtMkGO or the combined MitoTracker Green / TMRE) imaging used rapid switching of 475nm and 500-600nm LED modules with 474/27nm and 561/14nm clean-up excitation filters into secondary dual-band 482/563nm excitation filter, a 488/561nm dual-band dichroic, and a dual-band 523/610nm emission filter. Images were acquired with a Hamamatsu ORCA-Flash 4.0 CMOS camera controlled via PC with 3i Slidebook image capture software. Slidebook software integrated a six-channel perfusion system (VC6; Werner Instruments) and temperature control system via a 3i TTL breakout box. Temperature was maintained at 37°C with both in-line and platform heaters (Werner Instruments) with dual feedback circuits.

The same acquisition parameters were used across treatments. For TMRE/MitoTracker experiments, 1000ms were used for each channel. For mtMkGO experiments, the 475nm wavelength was acquired at 500ms and 600nm wavelength was acquired at 1000ms. For Mitophagy experiments, the 600 nm wavelength was acquired at 1000ms. For each experiment, the CamKII-constructs were acquired at 100ms.

For intraluminal lysosomal pH analysis, cells were transfected with hLAMP1Full-length-RpH-Lourin2. The imaging protocol used was set as following: 2 min in ACSF, 2 min in ACSF-NaProp 80 mM, 4 min in ACSF, 2 min in ACSF-NaCl 80 mM, 4 min in ACSF.

For mtMkGO analysis, cells were transfected mtMkGO construct at DD28 and analyzed at DD33.

For mitophagy analysis, cells were transfected with CamkII-GFP to identify single neurons. After 72h, iNeurons were treated with the Mitophagy Detection Kit (Dojindo Laboratories) following the manufacturer’s instructions. For mitochondria membrane potential analysis, cells were transfected with CamKII-Electra2 construct. After 72h, the cells were incubated with 10nM TMRE (Invitrogen) for 45 min and 50nM of MitoTracker Green (Invitrogen) for 15min.

For DQ-Red BSA analysis, iNeurons were transfected with CamKII-GFP for 72h and subsequently treated with 18 μg/ml DQ-Red BSA for 6 h. Coverslips were then washed twice with PBS and incubated with DAPI for 5min. After a PBS wash, the coverslips were mounted with ProLong Gold and imaged with Nikon TiE 63x-oil immersion after 16h.

For Acidifying Nanoparticle (AcNP) experiments, iNeurons were exposed to 1mg/mL of 647/680 fluorescent 100nm diameter nanoparticles (Degradex PLGA, Phosphorex, LGFR100). The AcNPs were added to the media for 72h, and cell were immediately after analyzed (3 days treatment). For long-term experiments, the iNeurons were exposed to AcNPs for 72h, the media was then replaced with BrainPhys for additional 3 days; the cells were then analyzed (6 days treatment).

## Image Analysis

Images were analyzed by an observer blind to experimental conditions using Fiji.

### Quantification of mitochondrial membrane potential

Rolling ball background subtraction (rolling ball radius = 50 pixels) was executed on both green and red images. MitoTracker Green is used to segment mitochondria in the corresponding CamKII-Electra2 positive cell. The ratio of the mean gray values for each segmented mitochondrion between the TMRE and MitoTracker signals is used to determine mitochondrial membrane polarization. The highest value for each image is reported to 1, and all the ROIs are compared to this one as a variation. Value for all the mitochondria are presented as a frequency distribution, setting the bin as 0.1

### Quantification of mitochondrial protein age

Rolling ball background subtraction (rolling ball radius = 50 pixels) was executed on both green and red images. A threshold was set to segment the mitochondria and identify ROIs. Ratio of the green/red mean gray value for each ROIs is used to determine the mitochondria age.

### Quantification of mitophagy

Rolling ball background subtraction (rolling ball radius = 40 pixels) was executed. A threshold was set to segment the mitochondria and identify ROIs. Red mean gray value for each ROIs is used to quantify mitophagy. The lowest mean gray value is set as the reference (=1) for each image, and all ROIs are reported relative to it. The mitophagy is presented as a frequency distribution plot with a bin set as 0.1.

### Colocalization between Lyso Dye and Mtphagy Dye

Colocalization was measured in Fiji by calculating the overlapping area between the Lyso Dye and Mitophagy Dye. Find Maxima function was used for Lyso Dye positive puncta, and auto-local threshold function with MidGrey method was used for Mtophagy Dye positive structures for each individual cell. The “and” function in the image calculator tool was used to measure the co-localization area.

### Quantification of lysosomal markers and DQ-BSA Red

Rolling ball background subtraction (rolling ball radius = 40 pixels) was executed. Threshold segmentation was used to identify and segment LAMP1-positive puncta or DQ-BSA puncta. The area and number of the identified puncta were measured using “analyze particles” setting the dimension at 20-200nm.

### Quantification of immunoblot

Protein levels were quantitated using the gel analyzer tool in ImageStudio. Boxes were used to designate the individual gel bands, and the integral of the intensity graph was taken as the value designating protein levels. ACTB is used as a loading control. Values were then normalized relative to the relevant control sample and expressed as fold effect.

### iPSC proteomics procedure

iPSC pellets were treated with acetone to concentrate and delipidate the proteins. Four volumes of-20°C analytical grade acetone (Sigma-Aldrich; St. Louis, MI) were added to each sample, mixed, and left overnight (6-8 h) at 4°C. The samples were centrifuged at 16,000 x g for 10 minutes at 4°C. After the supernatant was discarded, the pellet was washed twice with-20°C acetone and spun at 16,000 x g for 10 minutes at 4°. The pellet was then dried under a vacuum for 15 minutes on a SpeedVac. The pellets were solubilized in 100 μl of 6 M urea/50 mM ammonium bicarbonate, pH 8, and were subsequently treated with 2.5 μl of 5 mM dithiothreitol (DTT) and incubated for 30 minutes at 37°C. Twenty µl of 5 mM iodoacetamide (IAA) was added, and the solution was incubated for 30 minutes in the dark. Afterward, 20 μl of DTT was added and incubated for 10 minutes at 20-22°C to quench the IAA excess. A mass spectrometry grade mix of rLys-C and Trypsin Gold (Promega; Madison, WI) was added in a 1:25 ratio and incubated for 4 hours at 37°C. To dilute the urea concentration to >1 M, 600 μl of 50 mM ammonium bicarbonate was added and incubated overnight at 37°C. The following day, the digests were placed in a Macro Spin Column (The Nest Group, Inc., Ipswich, MA) to desalt. Around 10-100 μg of the digests were subjected to mass spectrometry analysis following the protocol described by (1). *Liquid Chromatography and Tandem Mass Spectroscopy* **-**The protein digests were randomized and analyzed using a Thermo Scientific Q Exactive Orbirtrap mass spectrometer fitted with a Thermo Scientific Proxeon Easy-nLC II HPLC and a Proxeon nanospray source at the Proteomics Facility at the University of California, Davis. The digests were loaded on a 100 μm x 25 mm Magic C18 200 Å 5U reverse phase trap where they were desalted online before being separated using a 75 μm x 150 mm Magic C18 200 Å 3U reverse phase column. The peptides were eluted using a 90-min gradient with a 300 nl/min flow rate. Am MS survey scan was obtained for the 300-1600 *m/z* range, MS/MS spectra were acquired where the top 15 ions in the MS spectra were subjected to high energy collisional dissociation, an isolation mass window of 2.0 *m/z* was used for precursor ion selection, and a 27% normalized collision energy for fragmentation with a 5-s duration for dynamic exclusion (1). *Protein Identification* **-** Using X! Tandem (THE GPM, thegpm.org; version X! Tandem Alanine (2017.2.1.4)) database searching all MS/MS samples to search the Felis catus database assuming digestion enzyme: trypsin, and searched with a fragment ion mass tolerance of 20 ppm and a parent ion tolerance of 20PPM, carbamidomethyl of cysteine, selenocysteine, ammonia-loss of the *N*-terminus, Gln->pyro-Glu of the *N*-terminus, deamidated of Asn and Gln, oxidation of Met and Trp and deoxidation of Met and Trp were all specified in X! Tandem as variable modifications. Weighted spectral counting was used to determine protein abundance (1). The criteria for protein identification-Scaffold (version Scaffold_4.11.1, Proteome Software Inc., Portland, OR) to validate MS/MS based peptide and protein identification were accepted if they could be established at greater than 88.0% probability to achieve a less than 0.5% FDR according to the Scaffold Local FDR algorithm. Acceptable protein identification was established at a probability greater than 5.0% to achieve an FDR less than 5.0%, and it contained at least 2 identifiable peptides, the probabilities of which were assigned using the Protein Prophet algorithm. Proteins with similar peptides not discernable with MS/MS were grouped to satisfy parsimony principles. *Protein profiling-*Features with >50% missing values were removed.

Missing values were imputed using the multivariate robust principal components, which replace missing values using a low-rank matrix factorization (singular value decomposition) robust to outliers (JMP Pro v17.0). The proteomes were filtered by unannotated features in the human protein database. The proteins were normalized by their sum, and log2-transformed and autoscaled (mean-centered and divided by the standard deviation of each variable). For the differential expression analysis, the primary metadata were the diagnoses; the reference group was the control; and all contrasts (ANOVA-style) were evaluated, followed by Fisher LSD post hoc. Statistically significant proteins were considered those with *p*-value < 0.05 with no cut-off for the fold change.

### RNA-seq statistical analysis

Raw counts were normalized using variance-stabilizing transformation and subsequently log-transformed to reduce heteroscedasticity and improve comparability across samples. To accommodate the complex experimental design and multiple metadata variables, we employed a multifactor linear modeling framework using limma with covariate adjustment, taking advantage of its high-performance empirical Bayes implementation for RNA-seq– derived expression values. Differential expression analyses were performed with diagnosis as the primary factor of interest, using either the control group or the non-deficient group as the designated reference. All pairwise contrasts were evaluated in an ANOVA-style design to identify diagnosis-dependent transcriptional changes across conditions. Statistical significance was determined using limma’s robust trend empirical Bayes moderation, applying a threshold of *p* < 0.05 without enforcing a fold-change cutoff, allowing unbiased detection of subtle but biologically meaningful differences in gene expression.

## Statistical Analyses

For proteomic datasets, protein abundance measurements were obtained in technical triplicates to ensure analytical reproducibility. For both proteomic and RNA-seq datasets, statistical significance was defined as *p* < 0.05 unless otherwise specified. To evaluate shifts in distributional profiles (e.g., cumulative frequency or density plots), the Kolmogorov–Smirnov test was applied, with significance determined at *p* < 0.05. For comparisons involving three or more experimental groups, we used one-way ANOVA followed by Tukey’s post-hoc test to correct for multiple pairwise comparisons and identify group-specific differences. All summary data are presented as mean ± standard deviation (SD) unless noted. Statistical analyses were performed using standard implementations in R or Python-based scientific computing packages.

## Results

### Schizophrenia in iNeurons from 22qDS is associated with older and less polarized mitochondria

Our previous study found that iNeurons from 22qDS +Sz had reduced ATP levels as well as Complex I and IV activity, while those from 22qDS-Sz had upregulated gene expression associated with mitochondrial biogenesis and mitophagy(*22*). Because both ATP synthesis and respiratory chain function are directly dependent on the mitochondrial membrane potential (ΔΨm)—the electrochemical gradient across the inner mitochondrial membrane that drives oxidative phosphorylation—alterations in ΔΨm represent a critical readout of overall mitochondrial health. Moreover, changes in biogenesis or turnover programs often manifest as shifts in ΔΨm, as newly generated or damaged mitochondria typically display distinct membrane potential profiles. Therefore, to determine whether the functional integrity of the mitochondrial network is preserved or compromised across groups, we quantified ΔΨm in iNeurons at differentiation day 30 (dd30) as a direct, sensitive indicator of mitochondrial competency and energetic status. Fluorescence signal of the positively charged dye tetramethylrhodamine ethyl ester (TMRE), normalized to the fluorescence signal from the biotracer MitoTracker Green, revealed a population shift towards reduced polarization in the 22qDS +Sz group compared to both the 22qDS-Sz and control groups (Figure 1A-1D; Figure S1A,B). Remarkably, the TMRE signal was significantly elevated in the 22qDS-Sz iNeurons relative to the controls, consistent with greater hyperpolarization. Of note, we performed all live-imaging experiments in iNeurons co-cultured on rat astrocytes, as astrocytes optimize the health of cultured neurons.

**Figure 1.**
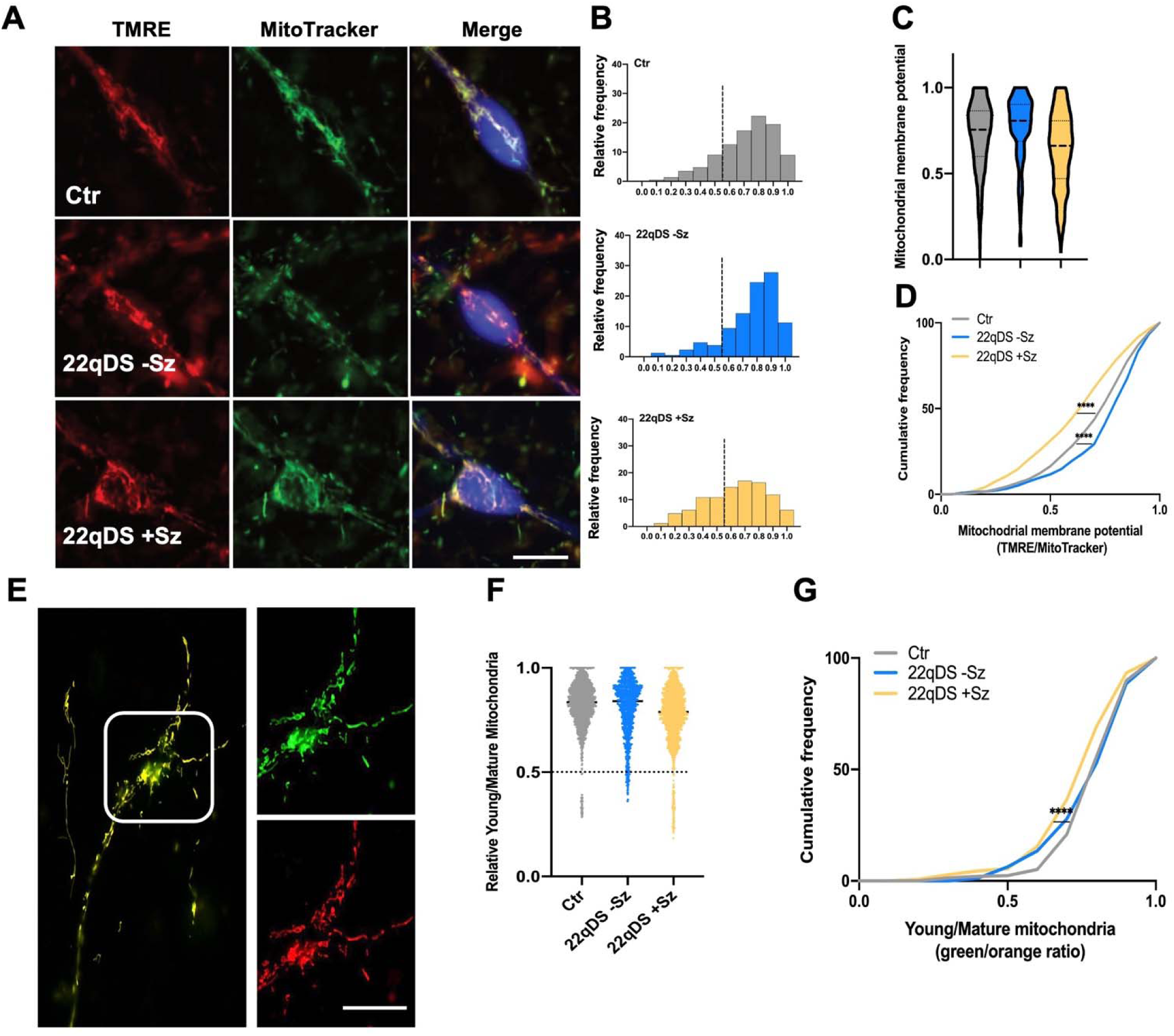
Schizophrenia in 22qDS is associated with a higher percentage of depolarized and aged mitochondria. **A)** TMRE (red) and MitoTracker (green) imaging of cells from control, 22qDS –Sz and 22qDS +Sz. Scale bar 50 µm. **B-D**) Quantification of TMRE relative to Mito Tracker fluorescence. **B)** Frequency distribution of relative mitochondrial membrane potential among the 3 groups. Dotted line separates the graph into relatively hyperpolarized versus relatively depolarized mito (<0.5). **C)** Violin plot and **D**) Cumulative distribution of the TMRE fluorescence signal normalized to the MitoTracker (mitos from n=6 lines for Ctr, n=5 lines for 22qDS-Sz and 22qDS +Sz). **E)** Representative images of control dd33 iNeurons transfected with mtMkGO. **F)** Quantification of fluorescence intensity of the ratio of young/old pool of mitochondria proteins in dd33 iNeurons **G)** Frequency distribution of mtMkGO signal. (mitos from n=6 lines for Ctr, n=5 lines for 22qDS-Sz and 22qDS +Sz). **** p<0.0001 Kolmogorov-Smirnov test between groups.

One mechanism for improving mitochondrial function in the context of genetic compromise is to enhance mitochondrial biogenesis and clearance of defective mitochondria(*34*). To determine whether these options may influence the penetrance of Sz in the context of 22qDS, we assess the mitochondrial turnover by using a modified timer protein MkGO(*35*) for optimal expression in human neurons. We fused this probe to mitochondrial localizing sequences of Cox8 (mtMkGO) to ensure mitochondrial targeting. mtMkGO achieves maximal fluorescence emission to green-wavelength excitation roughly 8 hours after its translation, whereas it achieves maximal fluorescence emission to red-wavelength excitation approximately 16 hours after its translation. Once steady state is achieved, in this case by transfection at dd28 and imaging 5 days later, the ratio of green to red emission is a relative measure of the flux of this construct and hence a relative measure of the turnover (or the balance between genesis and clearance of mitochondria). Indeed, treatment of iNeurons with the protein synthesis inhibitor anisomycin shifted the mtMkGO signal to “older”, whereas treatment with the PPARα agonist and activator of mitochondrial biogenesis bezafibrate shifted this signal to “younger” (Figure S1C). Comparison of mtMkGO signal in iNeurons from controls, 22qDS-Sz, and 22qDS +Sz revealed that the mitochondria population is relatively “younger” in the 22qDS-Sz group than in controls, but “older” in the 22qDS +Sz group than in controls (Figure 1E-G; Figure S1D), consistent with reduced mitochondrial turnover. Because mtDNA content did not differ across groups (Figure S1E), the observed changes in mitochondrial membrane potential likely reflect qualitative differences in mitochondrial function rather than differences in mitochondrial abundance. These findings indicate that mitochondrial health—as measured by inner membrane polarization—is impaired in 22qDS +Sz iNeurons.

They further suggest that mitochondrial biogenesis and turnover are reduced in this group relative to controls, whereas these processes appear to be upregulated in 22qDS-Sz iNeurons.

### Schizophrenia in 22qDS is associated with defective autophagy and mitophagy

These observations raise the question of why 22qDS +Sz iNeurons fail to mount the compensatory increases in mitochondrial biogenesis and turnover observed in the 22qDS-Sz group. The presence of a larger population of depolarized mitochondria in the +Sz group suggests that the removal of damaged organelles may be impaired. This pattern points toward a defect in autophagy—specifically mitophagy—as a potential mechanism underlying the inadequate turnover of dysfunctional mitochondria in 22qDS +Sz neurons.

To explore this possibility, we studied gene and protein expression in iNeurons differentiated for three weeks without astrocytes. mRNAs and proteins from iNeurons from n=4 iPSC lines in both 22qDS groups were normalized against the control group (also n=4 lines) and then compared to each other. From RNA analysis, upregulated pathways included “schizophrenia” as the strongest gene ontology (GO) term (Figure 2A).

**Figure 2.**
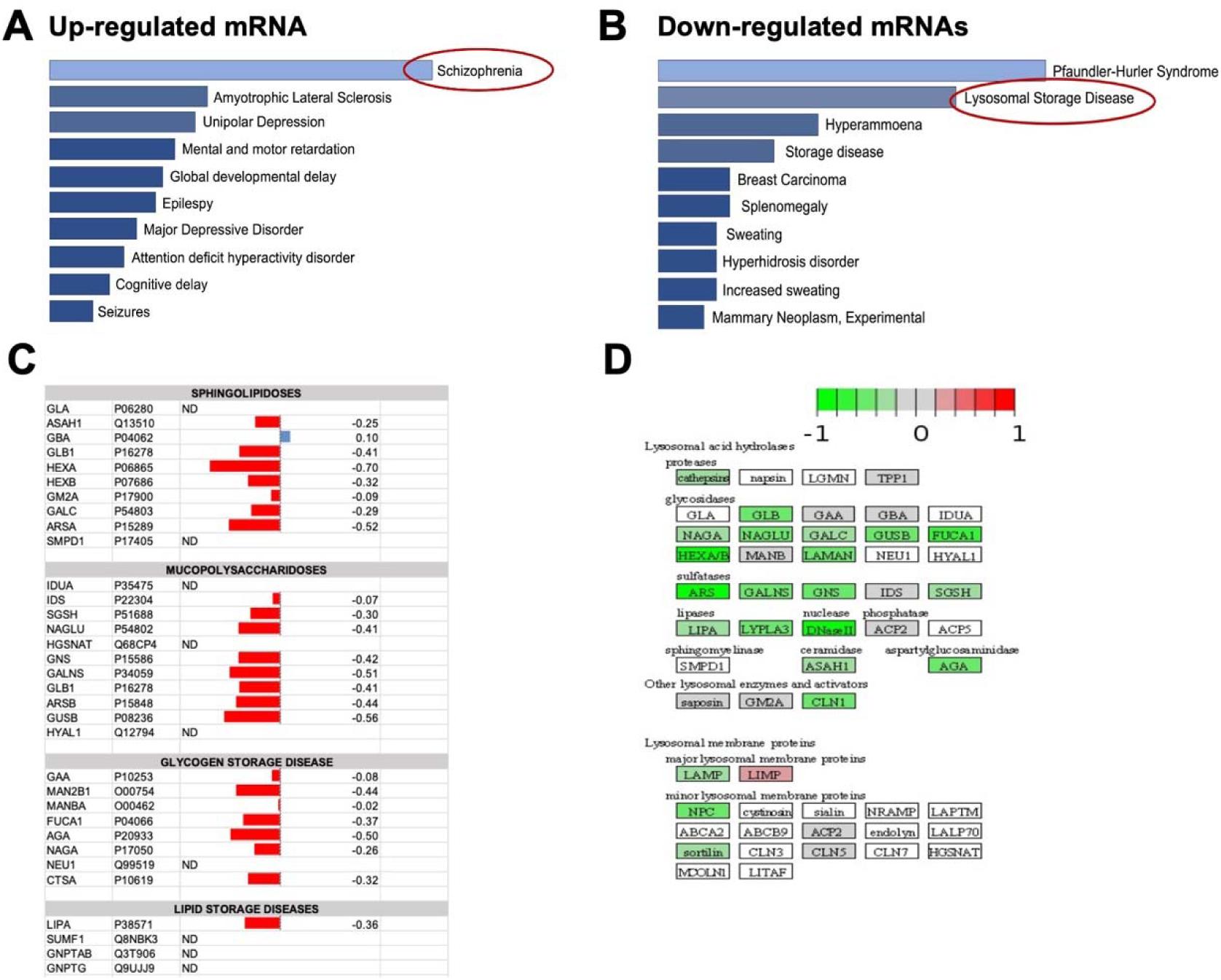
RNA sequencing and proteomics identifies lysosomal-related gene alterations in the 22qDS +Sz group. RNA and protein were harvested from dd21 iNeuron cultures from two runs of 4 iPSC lines each group (control, 22qDS –Sz and 22qDS +Sz, as per Li et al., 2021). The results are presented in terms of expression in the 22qDS groups relative to each other, normalized against those of the control group. **A-B**) Gene ontology (GO) pathway enrichment analysis identified (**A**) upregulated and (**B**) downregulated pathways in iNeurons from 22q +Sz compared to 22q –Sz. “Schizophrenia” is the most upregulated pathway in +Sz group, whereas “Lysosomal storage disease” pathway is strongly downregulated in the +Sz group. **C**) Expression level of proteins associated with lysosomal storage diseases again has a remarkable pattern of downregulation in 22qDS +Sz neurons compared to 22qDS –Sz neurons. **D**) Lysosomal proteins and enzymes showing downregulation in 22qDS +Sz compared to 22qDS –Sz. These results are consistent with lysosomal dysfunction in the 22qDS +Sz iNeurons.

Downregulated pathways in the 22qDS +Sz group included “Lysosomal Storage Disease” as second strongest GO term (Figure 2B). Selective analysis of the expression of individual proteins associated with lysosomal storage disease likewise showed a strong tendency for downregulation in the +Sz iNeurons (Figure 2C). Similarly, proteomic analyses also revealed a tendency for downregulation of lysosomal proteins and enzymes in the 22qDS +Sz iNeurons (Figure 2D).

Taken together, these results are consistent with reduced abundance of lysosome-associated proteins in the 22qDS +Sz group compared with both the 22qDS-Sz group and neurotypical controls, suggesting a potential deficit in lysosomal pathways. To further evaluate lysosomal status, iNeurons were again cultured to dd30 on astrocytes. As an initial structural measure, we assessed lysosome presence using immunofluorescence labeling for the lysosomal-associated membrane protein LAMP1. In the iNeuron somata from the 22qDS +Sz group there were reduced numbers of LAMP1 particles (Figure 3A,B), but these were of larger size (Figure 3A,C). We then employed the DQ-BSA assay in which red fluorescence emitted by particles bound to BSA becomes de-quenched by hydrolysis within the lysosome. The number of DQ-BSA particles is significantly reduced in the 22qDS +Sz iNeurons (Figure 3D,E). We also observed a trend for their size to be increased in the 22qDS +Sz group (Figure 3F). To examine one mechanism of reduced lysosomal function we then evaluated lysosomal intraluminal pH using a LAMP1-fusion of a pH-sensitive, ratiometric fluorescent protein (pHluorin)(*36*). Expression of this construct is driven by the neuronal-specific CamKII promoter and introduced by lipofectamine for sparse expression across the culture.

**Figure 3.**
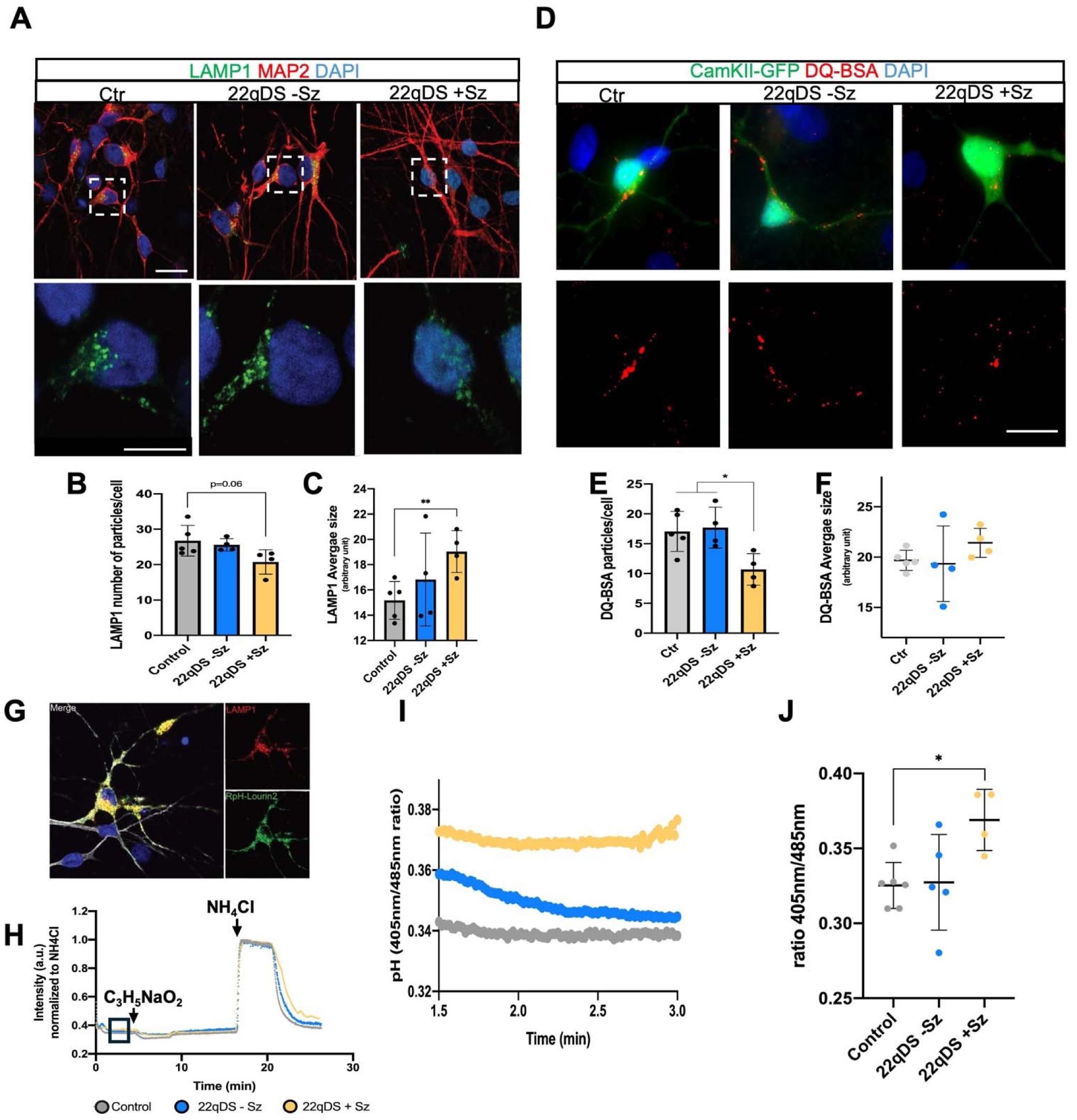
Lysosomal activity is reduced in 22qDS +Sz iNeurons. **A)** Representative images of dd30 iNeurons comparing 22qDS +Sz, 22qDS –Sz and healthy controls (Ctr) stained with the lysosomal marker LAMP1 and neuronal marker MAP2. **B-C**) Quantification of LAMP1+ particles/cell (**B**) and average size of the particles (**C**). Scale bar 50um and 200um. **D**) Representative images of iNeurons transfected with CamKII-GFP to analyze specifically human neurons at dd28. 48hrs after transfection the cells were incubated with 18ug/mL of DQ-BSA Red for 6h and then analyzed. Nuclei are stained with DAPI. Scale bar 50um. **E-F**) Number and size of active lysosomes of pre-loaded DQ-BSA as a readout of the lysosomal activity. **G**) Representative images of selective co-localization of the CamKII promoter-LAMP1-RpHLourin construct (green) and human LAMP1 (red) in iNeurons MAP2+ (blue). **H**) Representative trace of the fluorescence ratio of RpHlourin2 in response to acidification (Na+ Propionate) and alkalinization (NH4Cl) pulses in iPSC-derived neurons from healthy control patients and 22qDS patients with schizophrenia (22qDS +Sz) and without psychosis (22qDS-Sz). **I**) Highlight of the representative curves at resting conditions to clarify the differences in pH between samples. **J**) Quantification of the average pH at the timeframe of interest (as shown in Fig.5H), between different cell lines comparing 22q with and without Sz and neurotypical controls. Overall, these results show that lysosomes are dysfunctional in 22qDS +Sz iNeurons, with reduced degradative activity potentially due to elevated intraluminal pH. n=5 cell lines for Ctr, n=4 lines for 22qDS-Sz and 22qDS +Sz. * p<0.05 ** p<0.01 One-Way ANOVA, Tukey post-hoc test.

Specificity of the probe for human neurons was confirmed by selective co-labeling with human-specific LAMP1 (Figure 3G). Functionality of the probe was confirmed by response to acidification with sodium propionate and alkalinization with ammonium chloride (Figure 3H). Remarkably, lysosomal pH was elevated in the 22qDS +Sz neurons (Fig 3I-J).

The accumulation of relatively depolarized mitochondria in 22qDS +Sz iNeurons (Figure 1), combined with their reduced mitochondrial turnover and the lysosomal impairments identified in this group (Figures 2 and 3), suggests that schizophrenia in the context of 22qDS may involve a defect in mitophagy—the selective autophagic removal of damaged mitochondria. To assess mitophagy in iNeurons, dd30 cultures were exposed to a dye that selectively labels the mitochondrial outer membrane and fluoresces more intensely under low pH conditions (under pH 5; Dojindo), such as those found in lysosomes. However, while this Mtphagy dye indicates fusion of mitochondria to lysosomes, it does not indicate their actual degradation. Analysis was again restricted to neurons by reporter expression with the CamKII promoter (Figure 4A). Cumulative frequency distribution analysis revealed that relative to controls the mitochondrial localization to lysosomes was elevated in 22qDS +Sz iNeurons, whereas it was reduced in 22qDS-Sz iNeurons (Figure 4B–D). These findings align with our earlier observations of reduced lysosomal content, impaired proteolytic capacity, and elevated lysosomal pH in the +Sz group (Figures 2 and 3). Of note, Western blotting for proteins related to mitophagy and to autophagic flux, including LC3bII/l, were not as informative as the proteomic analysis due to high variability (Figure S2). However, optineurin levels were increased in the +Sz group, consistent with slowed autophagy/mitophagy (Figure S2). Together, these data suggest that less acidic lysosomes in 22qDS +Sz neurons slow mitophagy, causing mitochondria tagged for degradation to remain associated with lysosomes for longer periods. In contrast, lysosomal efficiency in 22qDS-Sz neurons appear to support more rapid turnover of damaged mitochondria, consistent with the elevated biogenesis and mitophagy gene expression previously observed.

**Figure 4.**
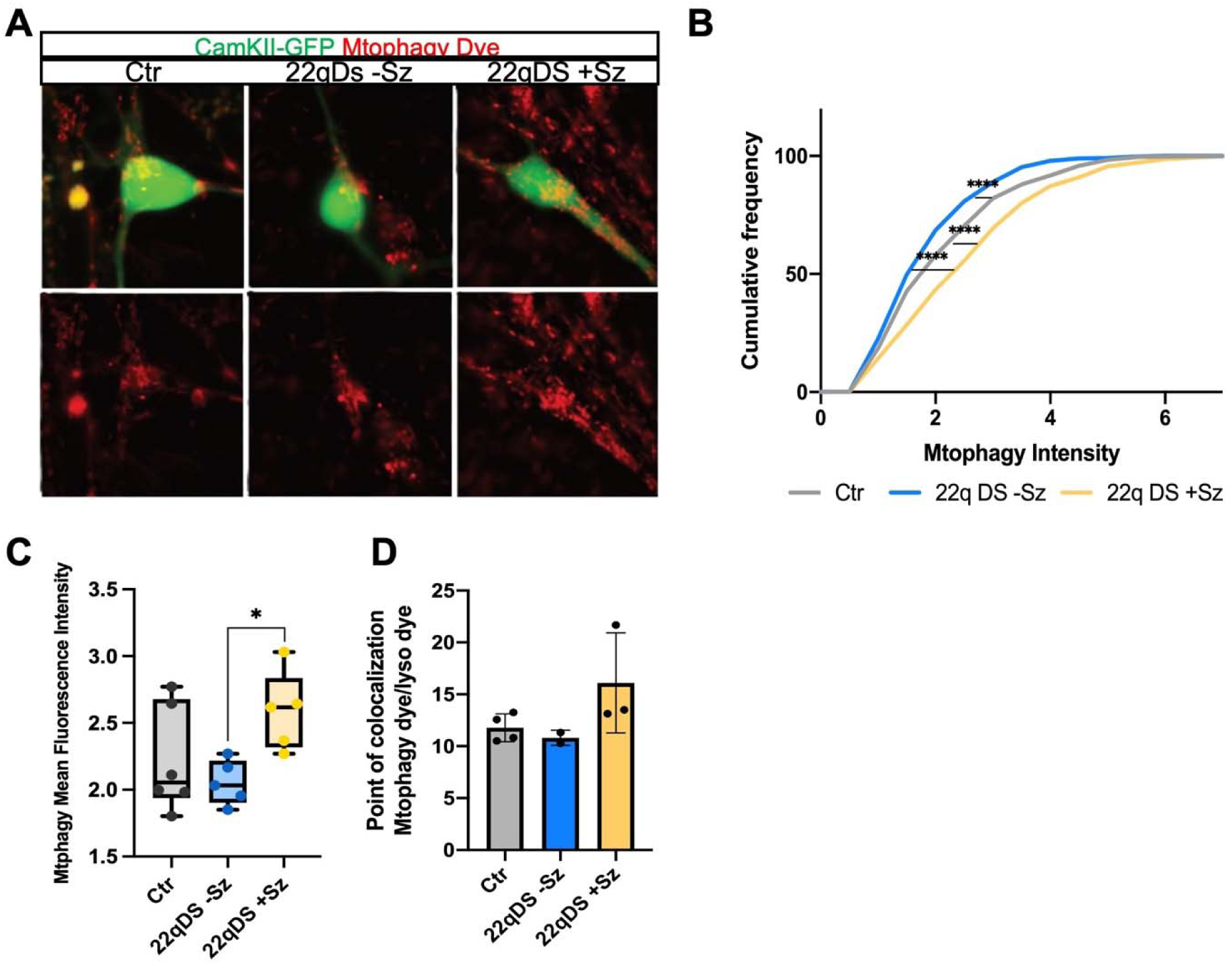
Alteration of mitophagy in the 22qDS +Sz iNeurons. Controls, 22qDS –Sz and 22qDS +Sz iNeurons were cultured on rat astrocytes. CamKII-GFP was introduced using lipofectamine at dd28 to sparsely label the iNeurons. Mitophagy was evaluated at dd30 with a mitochondria outer-membrane localizing, low-pH fluorescence-emitting dye (Mtphagy, Dojindo). The intensity of the dye increases when mitochondria are exposed to lysosomal acidification. **A**) Representative images of the mitophagy signal (red). For analysis this is normalized to the lowest signal of any mitochondria in that cell. **B**) Cumulative distribution of the mitophagy signal comparing the 3 groups, showing an increased signal in the 22qDS +Sz neurons. **C**) Mean fluorescence intensity analyzed across the lines. **D**) Co-localization of Mtphagy dye and Lyso dye (Dojindo), showing a trend for increased localization of mitochondria to lysosomes in 22qDS +Sz group. * p<0.05 One-Way ANOVA, Tukey post-hoc test. **** p<0.0001 Kolmogorov-Smirnov test between groups. n=6 lines for Controls, n=5 cell lines for 22qDS –Sz and 22qDS +Sz. For frequency distribution analysis, mitochondria from each line were pooled together.

### Acidifying nanoparticles restore lysosomal activity and enhance mitochondrial turnover in 22qDS +Sz iNeurons

Given that 22qDS +Sz iNeurons exhibit a modest level of lysosomal alkalinization, impaired degradative capacity, and slowed mitophagy, we tested whether restoring lysosomal acidification could rescue mitochondrial quality and degradation. We used poly-DL-lactide-co-glycolide (PLGA) acidifying nanoparticles (AcNPs) that enhance lysosomal acidification (*37, 38*)(Figure 5A). dd30 iNeurons were treated with AcNPs for 72 hours. AcNP treatment decreased lysosomal pH in 22qDS +Sz iNeurons to levels comparable with the 22qDS-Sz group (Figure 5B), and restored lysosomal degradative function as assessed by the DQ-BSA assay (Figure 5C,D). In line with these effects, AcNP-treated +Sz iNeurons displayed a modest but significant decrease in mitophagy signal compared with untreated +Sz cells (Figure 5E,F), suggesting that improved lysosomal acidification is accelerating the previously slowed clearance of damaged mitochondria.

**Figure 5.**
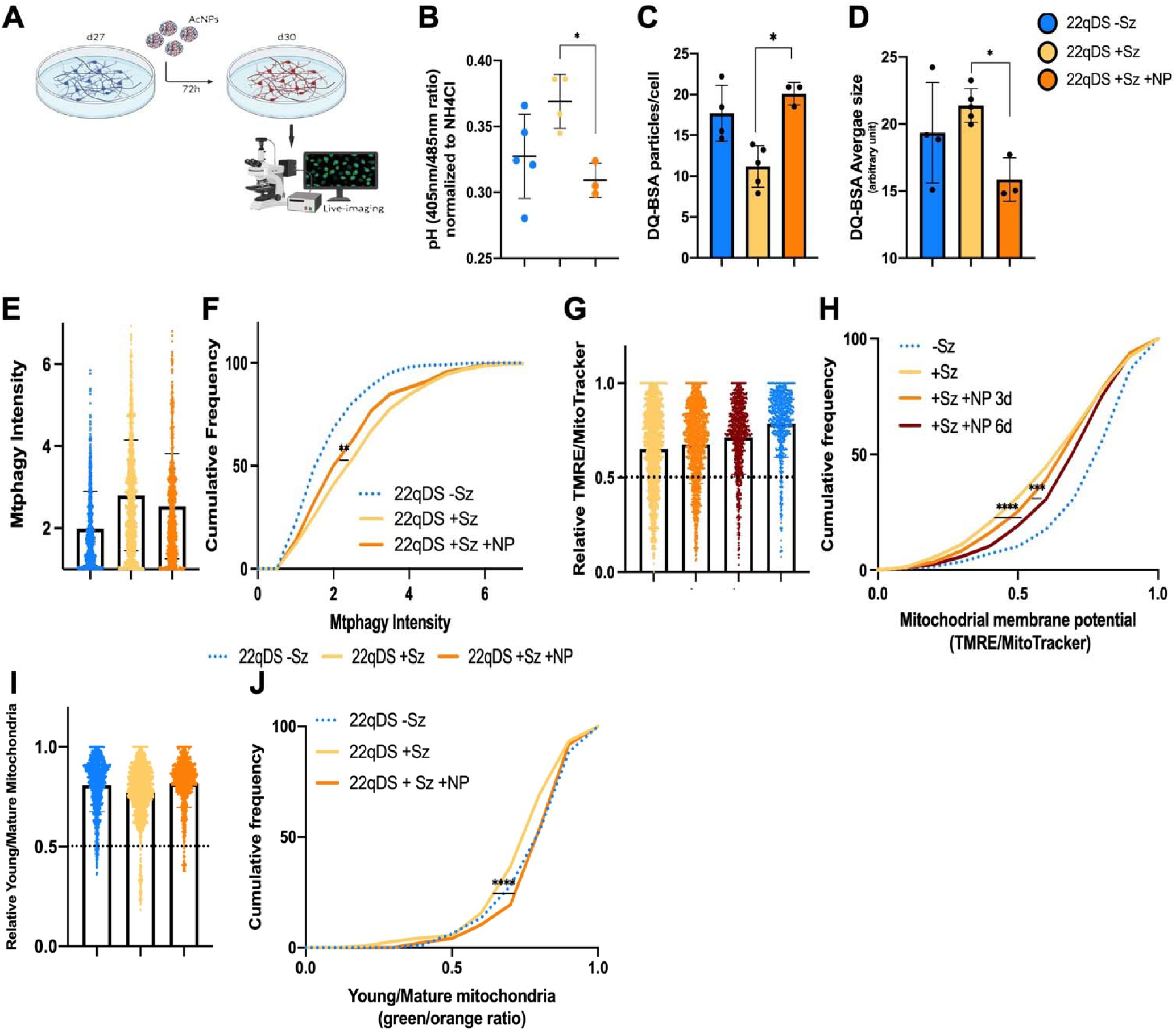
Acidifying nanoparticles (AcNPs) restore lysosomal activity in 22qDS +Sz iNeurons and improve mitochondrial health. **A)** Schematic of the use of AcNPs (1mg/mL for 72h) to acidify lysosomes. **B)** Quantification of the resting pH of 22qDS +Sz in response to AcNPs treatment compared to untreated 22qDS +Sz iNeurons. The values from the 22qDS-Sz group from Fig. 1 are used as reference control. **C-D**) Quantification of lysosomal activity in iNeurons transfected with CaMKII-GFP and incubated with AcNPs for 72h and with DQ-BSA for 6h. The number and size of DQ-BSA particles per cell are compared to untreated 22qDS +Sz and 22qDS –Sz iNeurons (from Fig.1). n=5 cel lines for 22qDS-Sz and 22qDS +Sz and n=3 cell lines for 22qDS +Sz +AcNPs. * p<0.05 One-Way ANOVA, Tukey post-hoc test. **E-F**) Localization of mitochondria to lysosomes is measured using the Mtphagy dye. the intensity of the dye is measured comparing 22qDS +Sz in the presence or absence of AcNPs. 22qDS –Sz is used as reference (blue dotted line) (from Figure 4). **G-H**) Mitochondrial membrane potential (and by extension mitochondrial health) is quantified using TMRE staining in the presence or absence of AcNPs in 22qDS +Sz iNeurons. AcNPs effects were analyzed after 3 days or 6 days. Cumulative frequency distribution of the ratio between TMRE/MitoTracker mitochondria is compared to 22qDS-Sz (from Fig. 1). **I-J**) Mitochondrial turnover is measured with mtMkGO in 22qDS +Sz in the presence or absence of AcNPs for 72h. 22qDS-Sz is used as reference control (from Fig. 1). ** p<0.01 ***p<0.001 **** p<0.0001 Kolmogorov-Smirnov test between groups. Mitochondria from n=6 lines for Controls, n=5 cell lines for 22qDS –Sz and 22qDS +Sz.

We next examined mitochondrial membrane potential using TMRE. Three days of AcNP treatment normalized the mitochondrial polarization of 22qDS +Sz iNeurons (Figure S3A-C), although these 22qDS +Sz neurons retained a higher proportion of hypopolarized mitochondria relative to treated-Sz cells (Figure 5G-H). Using mtMkGO to assess mitochondrial genesis and turnover, we also observed a reduction in the aged mitochondrial population in AcNP-treated +Sz iNeurons (Figure 5H,I). To determine whether more time may be needed for the treated lysosomes to clear depolarized mitochondria in the +Sz iNeurons, we examined TMRE signal again 3 days following washout of the AcNP treated media, and hence 6 days after the start of AcNP exposure. Indeed, this scenario resulted in further reduction of the relative population of depolarized mitochondria (Figure 5G-H).

Together, these results establish a mechanistic link between lysosomal alkalinization, slowed mitophagy, and impaired mitochondrial health in 22qDS +Sz iNeurons.

## Discussion

Previously, we found the iNeurons and LCLs from individuals with 22qDS +Sz had lower OXPHOS than 22qDS-Sz, while the 22qDS-Sz group had elevated expression of genes that control mitochondrial biogenesis and turnover(*22*). Here, we found that 22qDS +Sz IPSC-derived iNeurons had an increase of relatively depolarized mitochondria compared to both controls and 22qDS-Sz, (Figure 1). In contrast, iNeurons from the 22qDS-Sz group, compared to both controls and 22qDS +Sz, appeared to have younger mitochondrial proteins, consistent with the enhanced mitochondrial biogenesis implicated by gene expression data in our previous study and the active mitophagy (this study). The increased presence of relatively “aged” mitochondrial proteins and a relatively more depolarized population of mitochondria in +Sz iNeurons raised the question of whether mitophagy was altered in this group, a notion supported by RNA and proteomic evidence indicating lysosomal dysfunction (Figure 2).

Consistent with this notion, lysosomes in +Sz neurons were fewer in number but significantly enlarged, potentially reflecting the accumulation of undegraded cargo. This phenotype is of lower magnitude but reminiscent of observations in lysosomal storage disorders, such as Niemann-Pick disease, Gaucher disease, and Pompe disease, where impaired lysosomal enzyme activity leads to enlarged, dysfunctional lysosomes(*39, 40*). In the +Sz iNeurons, these enlarged lysosomes were also less functional, as shown by reduced DQ-BSA cleavage, and exhibited elevated intraluminal pH (Figure 3), indicating compromised degradative capacity.

Mitophagy assays revealed increased localization of mitochondria to lysosomes in 22qDS +Sz iNeurons (Figure 4). Together with the observation of enlarged lysosomes, these findings suggest that the elevated lysosomal pH in this group slows the degradation of depolarized mitochondria, leading to their accumulation. This interpretation is supported by experiments using acidifying nanoparticles (AcNPs; Figure 5). Treatment of 22qDS +Sz iNeurons with AcNPs restored lysosomal pH to levels comparable with controls and 22qDS-Sz neurons, and rescued lysosomal degradative capacity as assessed by the DQ-BSA assay. Importantly, this intervention also reduced the population of depolarized mitochondria, decreased mitochondrial localization to lysosomes, and improved mitochondrial protein “age,” indicating enhanced turnover. Together, these results suggest that schizophrenia in the context of 22qDS is associated with defective neuronal mitophagy, at least in part due to moderate lysosomal alkalinization, which impairs the clearance of damaged mitochondria and contributes to mitochondrial dysfunction.

The mechanism of elevated lysosomal pH is likely to include reduced ATP levels that we have previously shown to be present in 22qDS +Sz iNeurons(*22, 23*), since mitochondrial energetic defects can result in elevated lysosomal pH(*41*). This scenario raises a critical question: how does the process of becoming a compensator or non-compensator for ATP production via enhanced mitochondrial biogenesis unfold for a given cell, tissue, or person? As development proceeds for a cell type that requires oxidative phosphorylation for its function, such as neurons, a mismatch between ATP production and ATP requirement could develop in the context of 22qDS, in which 7 of the 45 affected genes encode mitochondrial-localized proteins. If the mitochondrial-nuclear signaling is adequate, then the nucleus could respond to the ATP mismatch by enhancing mitochondrial biogenesis and turnover. But if that signaling is relatively insensitive, such that mitochondrial ATP production falls to the point where lysosomal pH rises and mitophagy slows, the increasing population of relatively depolarized mitochondria would be expected to signal to the nucleus to reduce mitochondrial biogenesis(*42*). Potential contributors to reduced nuclear sensitivity include genetic variants outside the 22qDS locus that affect mitochondrial function, lysosomal regulation, or signaling pathways linking energy status to transcriptional programs. Environmental influences, such as oxidative stress, nutrient availability, or neuronal activity, may also modulate this compensatory response. Future studies will address these critical issues, but in addition to acidifying nanoparticles, there is tremendous interest in developing therapeutics for enhancing mitophagy in neurodegenerative disorders(*28, 43*), and diet and exercise can also have therapeutic effects via enhancement of mitophagy(*44*).

Outside of the 22qDS context, defective mitophagy has been implicated in both autism and schizophrenia. (*26, 30, 45*). How this affects brain function at the cellular level is unclear, although synaptic dysfunction appears to be one important component of both illnesses based on genetic and other studies(*46–49*), and mitophagy is an essential aspect of synaptogenesis, synaptic pruning, and synaptic function(*50–52*). While our study has the limitation of being based on a relatively small number of iPSC lines, in the context of previous studies on 22qDS and neurodevelopmental disorders, this study supports the contention that mitophagy may be a relatively underappreciated therapeutic target for schizophrenia and other genetic neurodevelopmental disorders in which mitochondrial dysfunction contributes to disease etiology.

## Acknowledgments

This work was supported by R01MH136706 (SA), NIH NS128751 (CG), DOD W81XWH2210560 (DCW), NIH CA259635 (DCW), NIH AG078814 (DCW).

## Author contributions

Conceptualization, S.A., C.G.; methodology, E.S., A.R.; investigation, E.S., A.R., J.L.; data curation, E.S., C.G., S.A.; formal analysis, E.S., S.A., C.G.; visualization, E.S. S.A.; writing—original draft, S.A., E.S.; writing—review and editing, E.S., S.A., C.G., D.W.; funding acquisition, S.A., C.G., D.W.; supervision, S.A., C.G..

