## Supplementary figures for "Variable penetrance of schizophrenia in 22q11.2 deletion syndrome is associated with impaired, but reversible, neuronal mitophagy"

**Supplementary Figures and Tables**

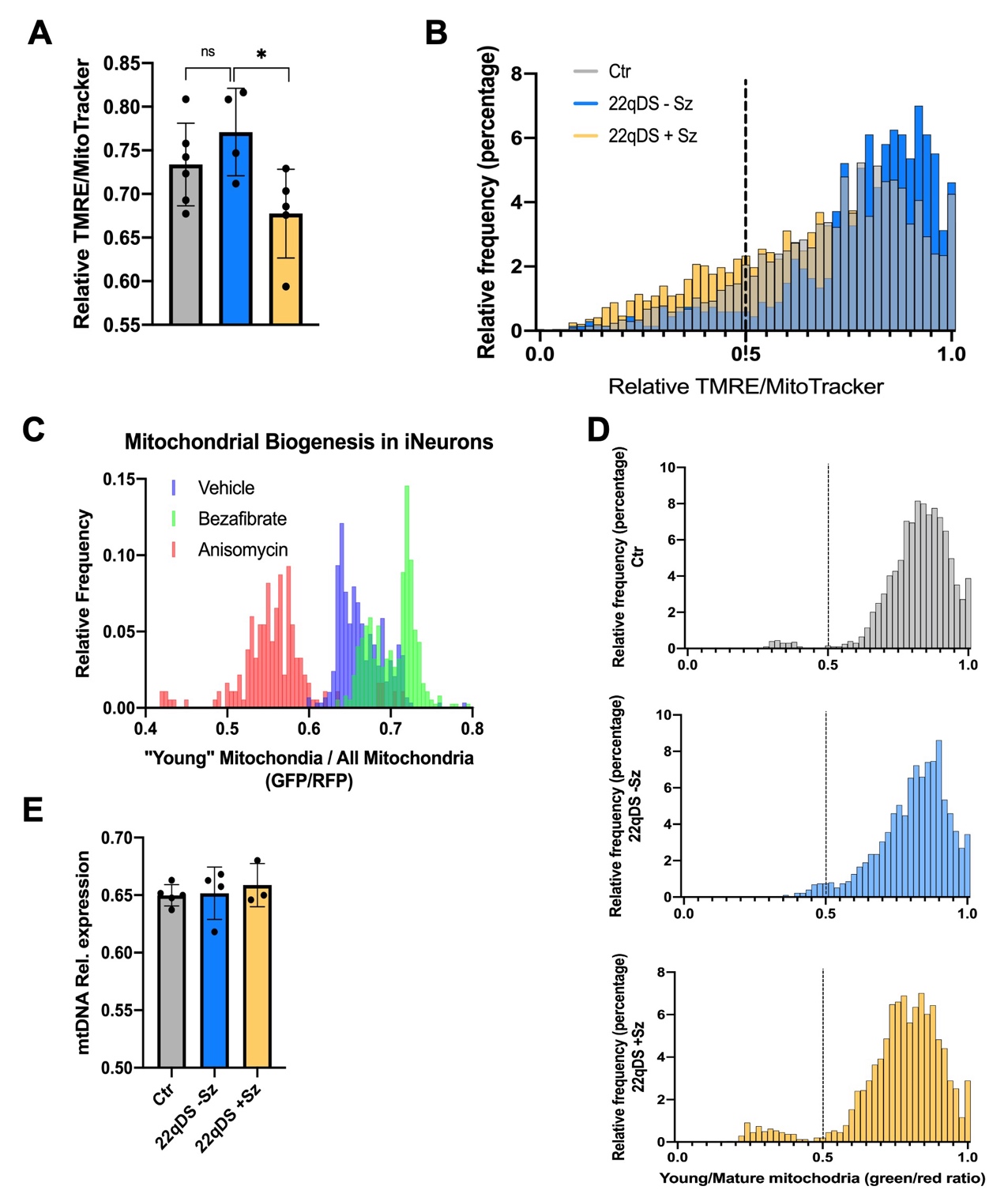

**Supplemental Figure 1. Schizophrenia in 22qDS is associated with higher proportion of depolarized and aged mitochondria**.

**A-B)** Quantification of fluorescence intensity of hyperpolarized mitochondria (stained by TMRE) compared to the total amount of mitochondria (stained with MitoTracker) in d30 iPSC-derive neurons. **A:** Average relative intensity of TMRE/MitoTracker between cell lines **B:** Frequency distribution of mitochondrial membrane potential among the 3 groups. Dotted line separates the graph into hyperpolarized mito (>0.5) and depolarized mito (<0.5)

**C)** Frequency distribution of green (younger) and red (older) mitochondria emission from d33 iNeurons from a control line transfected with mtMkGO. In presence of the protein synthesis inhibitor Anysomycin the green/red ratio shifts to the left (older), whereas in the presence of the PPARa agonist and driver of mitochondrial biogenesis Bezafibrate this ratio shifts to the right (younger).

**D)** Frequency distribution of relative mitochondrial age, as measured by mtMkGO, among the 3 groups. Dotted line separates the graph into relatively “younger” mito (<0.5) and “older” mito (>0.5). The 22qDS +Sz group shows an increased subset of older mitochondria protein pools compared to Ctr and 22qDS -Sz.

**E**) Mitochondrial DNA quantification comparing the 3 groups. *p<0.05 One-Way Anova, Tukey’s post-hoc test.

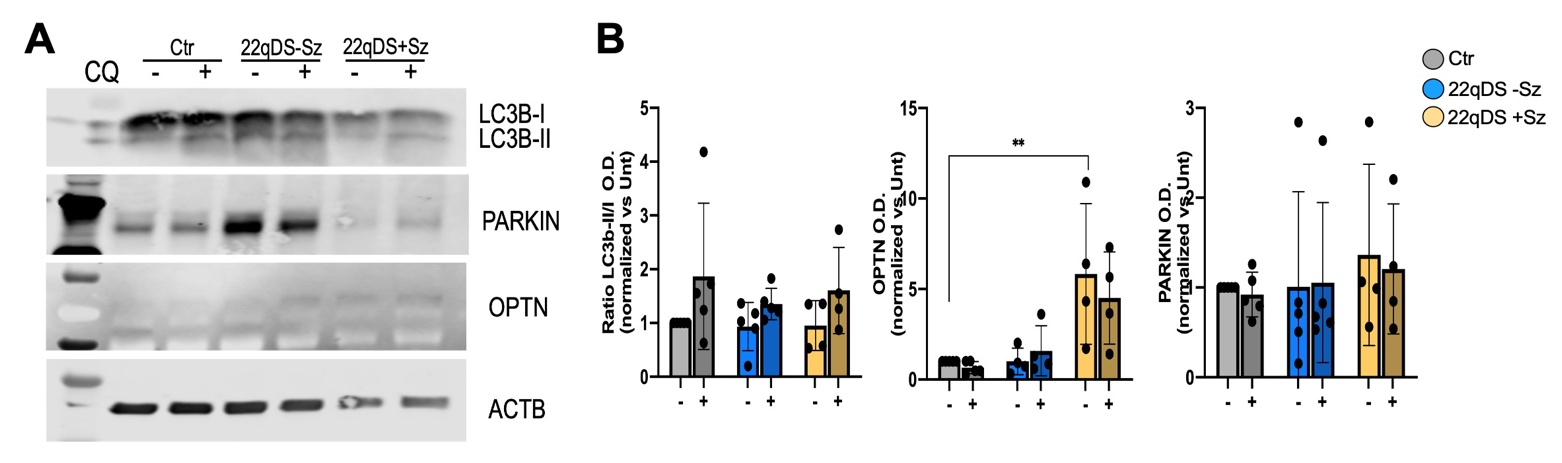

**Supplementary Figure 2: Expression of autophagy/mitophagy related transcripts and proteins.**

**A**) Representative western blot for autophagy markers and relative quantification (**B**) comparing the 3 groups. iNeurons were treated with Chloroquine (CQ) 1uM/6h to inhibit lysosomal degradation and evaluate mitophagy blockage. *p<0.05 **p<0.01 One-Way ANOVA test, Tukey’s post-hoc test. As evident in the dot plots, analysis of iNeuron differentiations by Western blotting was less informative due to high variability across differentiations. The imaging experiments may be more informative due to the presence of astrocytes that enhance the maturation and the electrophysiological activity of the iNeurons, and the use of CamKII promoter to drive reporter expression to identify the neurons for imaging also selects within the culture for more mature iNeurons.

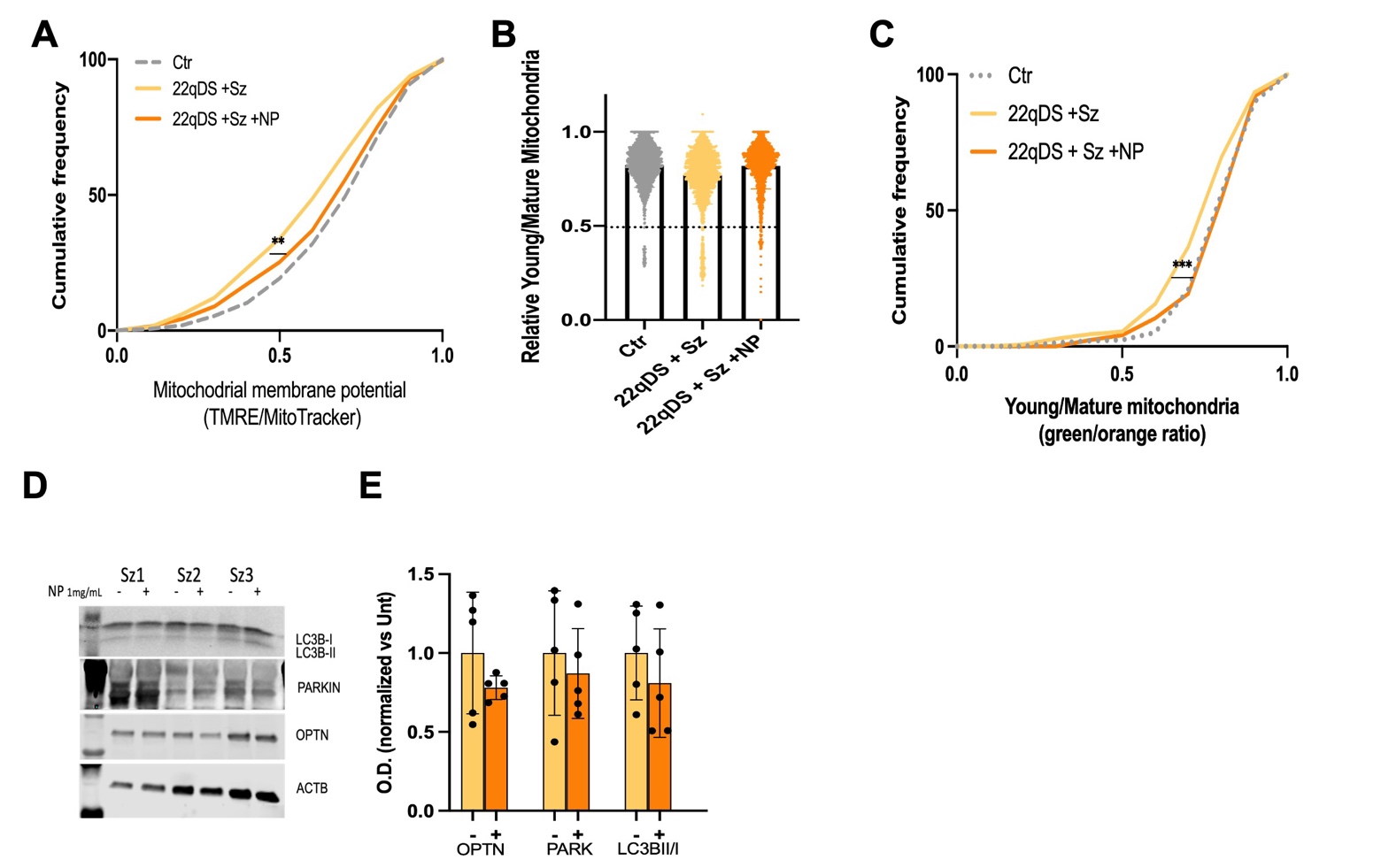

**Supplemental Figure 3. AcNPs restore mitochondrial membrane potential and maturation in 22qDS +Sz comparable to healthy controls**.

Compared to Figure 5, data on TMRE (**A**) and MkGO (**B-C**) in 22q +Sz iNeurons, treated with acidifying nanoparticles, relative to untreated 22qDS +Sz iNeurons and to control iNeurons (Figure 5H and 5J shows the comparison to untreated 22q -Sz iNeurons). **D-E**) Western blot of autophagy/mitophagy markers in 22qDS +Sz neurons treated with AcNPs 1mg/mL for 72h and relative quantification. Individual comparisons are not significant, but the trend for reduced optineurin, SQTMS1 (P62) and LC3Bll/l is consistent with the AcNPs improving lysosomal clearance of mitochondria and other organelles tagged for degradation.

**Supplementary Table S1**

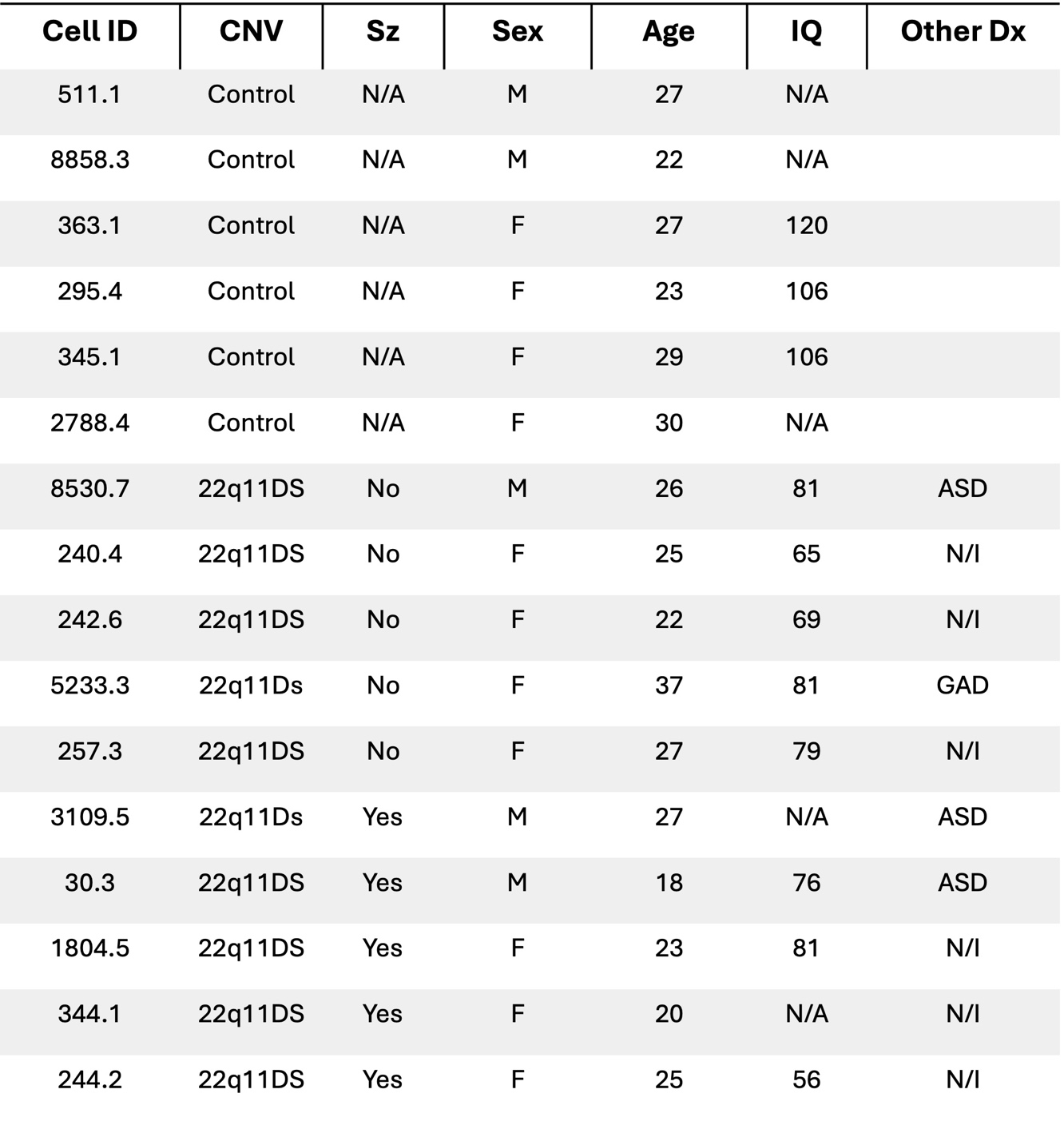

**Supplementary Table S1**. iPSC lines used in study. Lines were generated at Stanford University under the direction of Dr. Pasca, and the subjects were evaluated for the presence or absence of schizophrenia diagnosis. Information was obtained from the publication and from the NIMH Data Archive for R01MH100900. Numerical ID, sex and age of the subject at the time of biopsy collection, IQ and other comorbid psychiatry diagnoses are reported. Age refers to age when PBMCs were provided from which IPSC lines were generated. ASD Dx from ADOS, IQs from full-scale IQ-2. Average IQs controls: 110; 22q11DS –Sz: 75; 22q11DS +Sz: 71. IQ data was not included for two of the +Sz subjects, although it was reported that all +Sz subjects had verbal capacity adequate to assess psychosis. All lines were analyzed at CHOP for Multiplex Ligation-dependent Probe Amplification (MLPA) assay to verify sex and 22q11.2 deletion size. All subjects have the 3Mb “A-D” deletion, except 257.3 (-Sz) and 3109.5 (+Sz), who have the “A-C” deletion. N/I: not indicated; N/A: not available; ASD: autism spectrum disorder; GAD: generalized anxiety disorder.

**Supplementary Table S2**

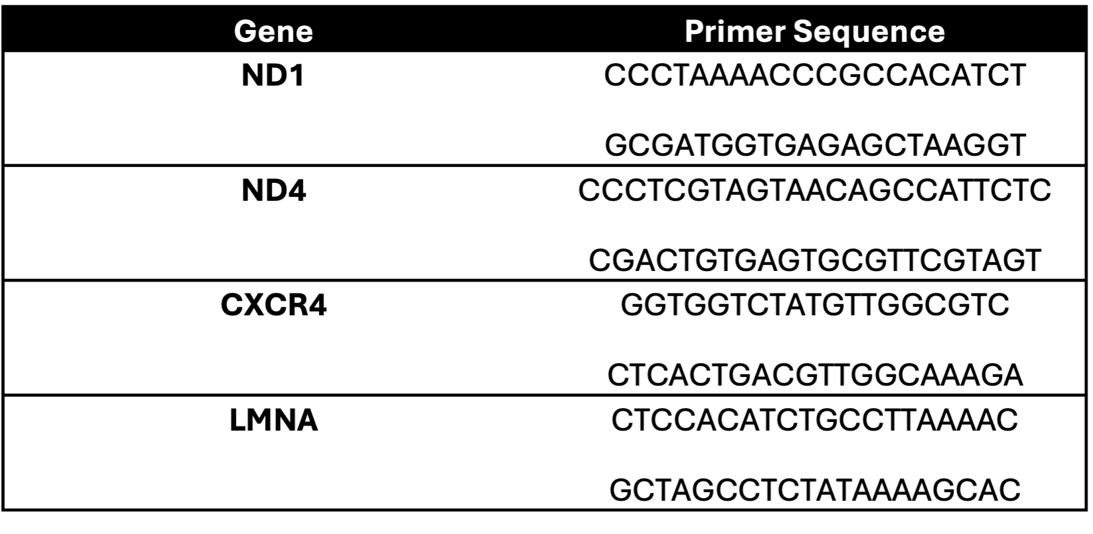
